# High-throughput nanopore sequencing of *Treponema pallidum* tandem repeat genes *arp* and *tp0470* reveals clade-specific patterns and recapitulates global whole genome phylogeny

**DOI:** 10.1101/2022.08.02.502389

**Authors:** Nicole AP Lieberman, Thaddeus D Armstrong, Benjamin Chung, Daniel Pfalmer, Christopher M Hennelly, Austin Haynes, Emily Romeis, Qian-Qiu Wang, Rui-Li Zhang, Cai-Xia Kou, Giulia Ciccarese, Ivano Dal Conte, Marco Cusini, Francesco Drago, Shu-ichi Nakayama, Kenichi Lee, Makoto Ohnishi, Kelika A Konda, Silver K Vargas, Maria Eguiluz, Carlos F Caceres, Jeffrey D Klausner, Oriol Mitja, Anne Rompalo, Fiona Mulcahy, Edward W Hook, Irving F Hoffmann, Mitch M Matoga, Heping Zheng, Bin Yang, Eduardo Lopez-Medina, Lady G Ramirez, Justin D Radolf, Kelly L Hawley, Juan C Salazar, Sheila A Lukehart, Arlene C Seña, Jonathan B Parr, Lorenzo Giacani, Alexander L Greninger

## Abstract

Sequencing of most *Treponema pallidum* (*T. pallidum*) genomes excludes repeat regions in *tp0470* and the *tp0433* gene, encoding the acidic repeat protein (*arp*). As a first step to understanding the evolution and function of these genes and the proteins they encode, we developed a protocol to nanopore sequence *tp0470* and *arp* genes from 212 clinical samples collected from ten countries on six continents. Both *tp0470* and *arp* repeat structures recapitulate the whole genome phylogeny, with subclade-specific patterns emerging. The number of *tp0470* repeats is on average appears to be higher in Nichols-like clade strains than in SS14-like clade strains. Consistent with previous studies, we found that 14-repeat *arp* sequences predominate across both major clades, but the combination and order of repeat type varies among subclades, with many *arp* sequence variants limited to a single subclade. Although strains that were closely related by whole genome sequencing frequently had the same *arp* repeat length, this was not always the case. Structural modelling of TP0470 suggested that the eight residue repeats form an extended α-helix, predicted to be periplasmic. Modeling of the ARP revealed a C-terminal sporulation-related repeat (SPOR) domain, predicted to bind denuded peptidoglycan, with repeat regions possibly incorporated into a highly charged β- sheet. Outside of the repeats, all TP0470 and ARP amino acid sequences were identical. Together, our data, along with functional considerations, suggests that both TP0470 and ARP proteins may be involved in *T. pallidum* cell envelope remodeling and homeostasis, with their highly plastic repeat regions playing as-yet-undetermined roles.

## Introduction

In recent years, efforts to catalog genomic diversity and phylodynamics of the syphilis spirochete, *Treponema pallidum* subsp. *pallidum* (*T. pallidum*), have resulted in a rapid increase in the amount of sequencing data and number of near-complete genome assemblies available in public databases (Arora et al., 2017; Beale et al., 2019, 2021; Chen et al., 2021; Grillová et al., 2019; Lieberman et al., 2021; Pinto et al., 2016; Taouk et al., 2022; Thurlow et al., 2022). Insights gained from these efforts, including the spread of azithromycin resistance (Beale et al., 2019) and high-resolution information on antigenic diversity (Lieberman et al., 2021), have aided our understanding of *T. pallidum* evolution and are invaluable to vaccine design. Although whole genome sequencing of low abundance *T. pallidum* DNA directly from clinical specimens is technically challenging due to the necessity of enrichment protocols such as hybrid capture with RNA or DNA baits (Pinto et al., 2016; Arora et al., 2017; Beale et al., 2019, 2021; Lieberman et al., 2021; Taouk et al., 2022), *Dpn1* enrichment (Grillová et al., 2019), whole genome amplification (Chen et al., 2021; Thurlow et al., 2022), and/or the traditional technique of passage of clinical strains through rabbits, sufficient progress has been made in development of these techniques that sequencing throughput of samples on a scale appropriate for monitoring of vaccine trials is feasible.

Most genomic analyses of *T. pallidum* have excluded portions of the genome difficult to resolve by short-read sequencing, including the *<u>T</u>. <u>p</u>allidum* <u>r</u>epeat (*tpr*) family of paralogous genes, the number of 60 bp tandem near-perfect repeats in the gene encoding the acidic repeat protein (*arp; tp0433*), and the number of 24 bp tandem repeats in the gene encoding the tetratricopeptide repeat protein TP0470 (*tp0470*). *Arp* repeat length has been interrogated extensively using the CDC (Pillay et al., 1998) and enhanced CDC (Marra et al., 2010) typing schemes. This has allowed monitoring of local strain composition over time (Marra et al., 2010; Flasarová et al., 2012; Grimes et al., 2012) and of strains circulating worldwide (Sutton et al., 2001; Pillay et al., 2002; Pope et al., 2005; Liu et al., 2020), and identification of subtypes enriched in neurosyphilis (Molepo et al., 2007; Marra et al., 2010). However, these approaches focus on obtaining the number of *arp* repeats, rather than providing sequence information on the repeats, which is necessary for more robust genotyping. Our sequencing-based approach starts to fill this important knowledge gap.

Little is known about the role of *tp0470* in syphilis pathogenesis, though tetratricopeptide repeat proteins in other bacterial species act as scaffolds for protein-protein interactions and are often critical to functionality of virulence factors (Cerveny et al., 2013). The highly charged motif “EAEEARRK”, encoded by the 24 bp repeat in *tp0470*, occurs C-terminal to the predicted tetratricopeptide repeat domain. This motif is repeated between 4-29 times in publicly available *T. pallidum* subsp. *pallidum* complete genomes, and up to 37 times in the *T. pallidum* subsp. *pertenue* strain CDC-2. The role of the *arp* gene is similarly understudied. It encodes an antigenic (Liu et al., 2007) protein containing at least four types of the 60 bp repeat that have been previously identified in *T. pallidum* subsp. *pallidum*, with differences confined to six positions, all of which result in amino acid substitutions (Liu et al., 2007; Harper et al., 2008). Repeat lengths between 2-22 have been previously observed in *T. pallidum* subsp. *pallidum* clinical specimens (Harper et al., 2008). Although the role of the acidic repeat protein in pathogenesis and colonization of various anatomic sites is unknown, some samples collected from whole blood had a lower number of *arp* tandem repeats than in patient-matched lesion swabs (Mikalová et al., 2013), and late stage syphilis samples are reported to have fewer *arp* repeats on average than early stage (Harper et al., 2008). Importantly, *arp* and *tp0470* are thought to evolve via intra-strain recombination (Grillová et al., 2019; Noda et al., 2022).

To date, most closed *T. pallidum* genomes have relied on Sanger sequencing of *arp* and *tp0470* (Cejková et al., 2012; Pětrošová et al., 2012; Zobaníková et al., 2013; Grillová et al., 2019), requiring extensive manual curation of data as well as ample starting material. Herein, we describe new bench and bioinformatic protocols for highly multiplexed nanopore sequencing of *arp* and *tp0470*, reducing the quantity of sample needed, as well as per-sample cost and hands-on time. These methodological improvements allowed us to gain insight into the evolution of both *tp0470* and *arp* and formulate hypotheses that pave the way for functional studies to understand the role of these putative virulence factors in syphilis pathogenesis.

## Methods

### Ethics Statement

All human samples were collected and deidentified following protocols established at each institution. All IRB information from samples collected in Japan; Italy; Ireland; Maryland, USA; Madagascar; Peru; Papua New Guinea, and Nanjing, China has been previously published (Hopkins et al., 2004; Lukehart et al., 2004; Van Damme et al., 2009; Hook III et al., 2010; Marra et al., 2010; Lieberman et al., 2021). Collection of additional samples was covered by the following IRBs: Malawi: National Health Sciences Research Committee Ministry of Health and Population (IRB Approval Number 2252); Colombia: Centro Internacional de Entrenamiento e Investigaciones Medicas (CIDEIM) Institutional Human Research Ethics Committee (CIEIH) (IRB protocol number 1289); Guangzhou, China: Dermatology Hospital of Southern Medical University (SMU) Medical Ethic Committee (IRB protocol number GDDHLS-20181202(R3)); Chapel Hill, USA: University of North Carolina IRB Protocol Number 19-0311. Sequencing of deidentified strains was covered by the University of Washington Institutional Review Board (IRB) protocol number STUDY00000885 and University of North Carolina IRB protocol number 19-0311.

### Whole genome sequencing and maximum likelihood phylogeny

Samples were sequenced and genomes assembled as previously described (Lieberman et al., 2021) or, for samples from Colombia, Malawi, Guangzhou, China, and Chapel Hill, USA, consensus sequences were assembled as previously described with minor modifications (Chen et al., 2021; Parr, 2021). Alignment, recombination masking and generation of the maximum likelihood phylogeny were performed as previously described (Lieberman et al., 2021).

### Barcoding PCR of *arp* and *tp0470*

Repeat regions of *arp* and *tp0470* were amplified using 96 combinations of forward and reverse barcodes barcoded primers containing a 24bp index (Supporting Information). Input volume ranged between 0.5-8 µL depending on genome copy number and amount of sample available. Samples were amplified by one of two methods, both using the Takara PrimeSTAR GXL polymerase in a 25 µL reaction: Those with low volume were first amplified with non-barcoded primers (98°C for 2 minutes, 35 cycles of 98°C for 10 seconds, 62°C for 15 seconds, 68°C for 2 minutes, then held at 68°C for 10 minutes before storing at 4°C), the amplicon cleaned with 0.8x Ampure XP beads, diluted 1:100, and then 1 µL template barcoded with 14 additional cycles of PCR using the indexed primers, using a 65°C annealing temperature. Alternatively, samples were amplified directly from the genomic DNA using barcoded primers. Both methods produced equivalent results and technical replicates for each method agreed with each other (Supporting Information, “Methods Equivalency”)).

PCR products were electrophoresed using 1% or 2% TAE agarose gels and purified with 0.6x or 0.8x volumes of AMPure XP beads for *arp* and *tp0470* amplicons, respectively. Clean PCR products were quantified using Qubit 1X dsDNA HS buffer (Invitrogen).

### Nanopore library preparation and sequencing

Following barcoding PCR and purification/quantification, samples were pooled to meet recommended input amounts for Oxford Nanopore (ONT) Adapter ligation kit (SQK-LSK109). Because most samples fell in the range of 2-10 ng/uL, we chose to maximize efficiency by pooling an equivalent volume (0.5 µL) per amplicon.

DNA end repair of the pool was performed as outlined in the ONT protocol for SQK-LSK109 using both ONT and NEBNext reagents, then purified with Ampure XP beads at a 1:1 ratio. Adapter ligation proceeded according to the manufacturer’s protocol, with room temperature incubation for 10 minutes, followed by purification with a ratio of 0.8x Ampure XP beads, washing with kit short fragment buffer (SFB), and elution in 47 µL water. The resulting pool was then quantified on Qubit (1X high sensitivity dsDNA kit) to calculate the total molecular weight of DNA. Molarity was calculated assuming an average fragment size of 800bp.

Amplicons were sequenced on Flongle flow cells via the MinION mk1B platform. ONT MinKNOW software interfaced with the MinION to perform pre-run flow cell checks and initiate/monitor the sequencing experiment. Each Flongle was primed and loaded with 20 fmol DNA per SQK-LSK109 and Flongle sequencing expansion (EXP-FSE001).

Sequencing was run for 24 hours, selecting “SQK-LSK109” for the DNA amplicon kit with high accuracy basecalling and other default parameters. Basecalling was performed in MinKNOW (v21.11.7) running Guppy (v.5.1.12) (Wick et al., 2019). Reads with an average phred score greater than 9 passed the quality filter.

### Demultiplexing

Fastq files that passed the quality filter were processed through Porechop (Wick et al., 2017) twice: First, using a customized adapters.py script that contained forward barcodes, using the default stringency of up to 5 mismatches in the 24 bp barcodes. Barcodes were modeled after those used previously in a dual-indexing protocol (Currin et al., 2019). Second round demultiplexing proceeded using reverse primers with the same parameters in the adapters2.py script, but ensuring no additional bases were trimmed past the barcodes.

### *arp* and *tp0470* consensus generation

Demultiplexed reads were aligned with bwa mem 0.7.17 (Li and Durbin, 2009) using the nanopore preset to reference files containing various numbers of tandem *arp* or *tp0470* repeats and flanking regions of several hundred bases. Per-reference statistics were extracted with the BBtools v38.18 (Bushnell) utility pileup.sh, and the reference assigned the most reads determined for each sample using a custom R script. Consensus sequences were extracted using the sam2consensus utility script (https://github.com/edgardomortiz/sam2consensus). Predicted number of repeats and resultant band size was then cross referenced with the agarose gel images, and the automated call either confirmed or overridden.

### Structural modeling

Locations of predicted signal peptides and lipidation sites were determined in “slow” mode in SignalP 6.0 (Teufel et al., 2022). Default settings were used for PSORT analysis (Yu et al., 2010). Conserved domains were determined using the NCBI Conserved Domain Database (Lu et al., 2020). Following removal of predicted secretion sequences, full length protein sequences for TP0470 and ARP were modeled in trRosetta (Yang et al., 2020; Du et al., 2021) with default settings and AlphaFold (Jumper et al., 2021; Varadi et al., 2022) using the pdb70 database for homology modeling and otherwise default settings. Protein models were visualized in PyMOL, and electrostatic surface potential shown with the Adaptive Poisson-Boltzmann Solver (APBS) plugin.

### Statistics and code availability

All statistical analysis was performed in R v4.0.3. Phylogenetic trees and *tp0470* and *arp* variants were visualized with the R packages ggtree (Yu et al., 2017), treeio (Wang et al., 2020), and ggplot (Wickham, 2016), and multiple sequence alignments by R package ggmsa (Zhou et al., 2022). Bash, R, and python scripts for all data processing are available at https://github.com/greninger-lab/TP_genome_finishing.

## Results

### Maximum likelihood phylogeny of 207 *T. pallidum* subsp. *pallidum* strains

We have previously reported the near-complete genomes of 196 *T. pallidum* strains, of which 191 were *T. pallidum* subsp. *pallidum* (Lieberman et al., 2021). Herein, we have updated the *T. pallidum* subsp. *pallidum* whole genome phylogeny to include 16 additional strains collected in Colombia, China, Malawi, and the United States. With the exception of one strain collected from Malawi (TPVMW082H), all newly-added strains fit within the subclades defined previously (Lieberman et al., 2021) (Fig 1A; Supporting Information). Malawi strain TPVMW082H appears to have diverged from the lineage that gave rise to the Nichols B and Nichols C subclades, but the maximum likelihood support values were below 0.95; therefore the phylogenetic relationship between TPVMW082H and Nichols B and C could not be clearly delineated. To demonstrate the fact that strain TPVMW082H is only distantly related to other samples included in this phylogeny, we have assigned this strain its own subclade, Nichols X.

**Figure 1:**
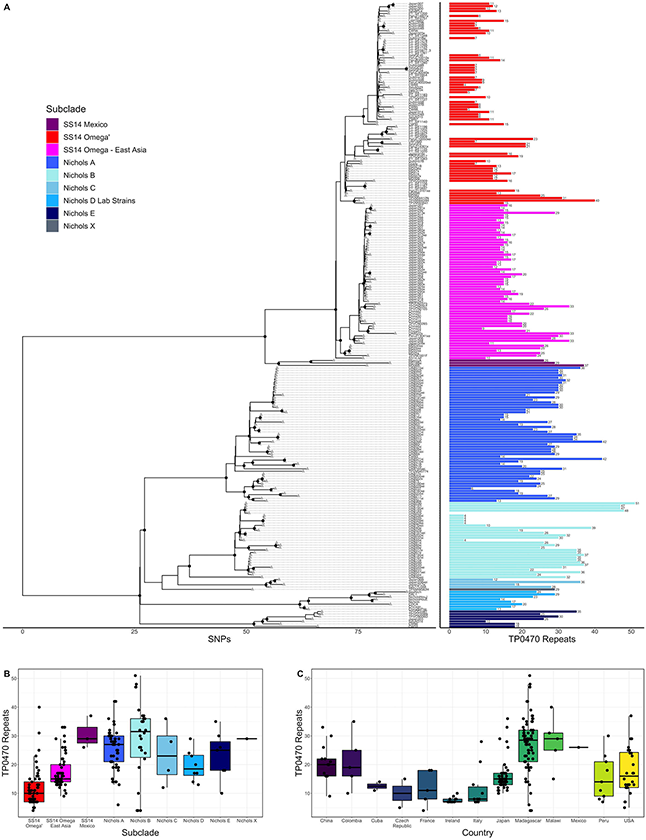
Variation in *tp0470* repeat length. A) Recombination masked whole genome phylogeny (left) with the number of *tp0470* repeats for each strain (right). Sequence variant number is included as text to the right of each length bar. All data are also included in a tabular representation in Supporting Info. Number of *tp0470* repeats by subclade (B) or country (C).

### Nanopore sequencing can resolve *tp0470* and *arp* repeat sequences in a high throughput fashion

To date, no systematic analysis of full-length *T. pallidum* tandem repeat genes has been performed, in part due to the fact that the short-read Illumina data used to generate most whole genomes cannot resolve the many 24- and 60-bp repeats found in *tp0470* and *arp*, respectively. Although Sanger sequencing can be employed to examine repeats from both genes, this approach is labor intensive and relies on the *a priori* assumption that the tandem repeats comprise fewer than ∼800 bp, the length limit for Sanger sequencing; lengths of up to 22 repeats (1320 bases) have been reported for *arp* (Harper et al., 2008). For *tp0470*, the DAL1 strain has 29 repeats of 24 bases (696 bases total) (Cejková et al., 2012), and the *T. pallidum* subsp. *pertenue* strain CDC-2 has 37 24 base repeats (888 bases total) with no upper limit known. Therefore, we developed bench and bioinformatic protocols for highly multiplexed, long read sequencing of the *arp* and *tp0470* loci. We amplified portions of the *arp* and *tp0470* genes using dual-indexed primers, allowing up to 192 amplicons on a single nanopore Flongle, followed by demultiplexing, reference mapping, and consensus calling. Across six Flongle runs to generate *arp* and *tp0470* data on 212 strains, an average of 408,286 reads passing filter were generated. Following demultiplexing of forward and reverse barcodes, an average of 3327 reads were assigned to each sample, comprising both *arp* and *tp0470* reads. Reads were then mapped to *tp0470* and *arp* reference files containing between 1-60 and 1-24 tandem repeats, respectively, using bwa mem (Li and Durbin, 2009) with the Oxford Nanopore preset (-ont2d) to account for low fidelity reads.

To validate our method, we cross referenced the band size seen on gel electrophoresis with the automated call from our pipeline. S1 and S2 Figs show the gel electrophoresis bands from a subset of samples and histograms of the distribution of mapping to the number of repeats for *tp0470* and *arp*, respectively. We found 89% (183/206) concordance between the *tp0470* pipeline call and band size, with discordance likely due to considerable low molecular weight byproducts produced during amplification of *tp0470* repeats, which are 75% GC-rich. However, as is clear for sample China11, the correct repeat number was readily apparent in these samples upon inspection of the mapping distribution (boxed region and arrows, S1B Fig).

Technical replicates of select samples agreed with each other in 100% of cases, however, amplicons produced using a two-stage amplification (see Methods) required more manual comparison with band size to eliminate the low molecular weight byproducts, which were unsurprisingly more likely to appear during two stage barcoding. Replicates generated using both methods gave the same result (Supporting Information, “Methods Equivalency”).

At 98.5% (200/203), concordance between the number of *arp* repeats determined automatically by our pipeline and the electrophoresis band size was extremely high (S2 Fig). Sanger sequencing was used to validate *tp0470* repeat lengths of select samples, with 100% concordance seen. Confirmation of novel *arp* repeat types, as well as linkage between adjacent 60 bp repeats, was performed by manual inspection of Illumina WGS reads for all unique *arp* variants. Technical replicates and replicates using each barcoding method gave the same result.

### The number of *tp0470* repeats is higher in samples from the Nichols-like clade

We first examined the consensus sequence and number of tandem 24 bp repeats in the *tp0470* gene in the context of the recombination-masked whole genome phylogeny we had previously determined (Lieberman et al., 2021). Unlike *arp*, which has some sequence variation at six positions per 60 bp repeat (see below; (Liu et al., 2007; Harper et al., 2008)), we did not find any *tp0470* sequence changes in any of the 233 strains we examined, which included 211 strains we successfully sequenced by nanopore and the remainder from public databases. However, we did note wide variability in the number of repeats (Fig 1A), ranging between 4-51, with Nichols-like strains generally having more repeats than SS14-like strains (mean (median) repeats for Nichols and SS14 25.7 (27) and 15.3 (14), respectively; *p*<2.2×10^−16^, Welch’s t-test) (Fig 1B), although this effect was difficult to disentangle from the strong location-specific effects we observed (Fig 1C, *p*<2×10^−16^, ANOVA).

### The number of *arp* repeats varies per subclade

We then examined the number of *arp* repeats in each sample in the whole genome phylogeny. Among the 226 TPA strains with *arp* sequence information, including 203 strains we sequenced and the remainder from public databases, the majority (170, 83.7%) had 14 tandem repeats, 28 (13.8%) had ten repeats, and the remainder had between 4-24 repeats (Figure 2A, Supporting Information), consistent with previous reports of the prevalence of repeat lengths (Pillay et al., 1998; Harper et al., 2008; Marra et al., 2010). No relationship between *tp0470* repeat length and *arp* repeat length was observed (SFig3A, Pearson coefficient = 0.008), as seen in prior observations (Šmajs et al., 2018), even when strains containing 14 repeat *arp* sequences were removed (Pearson coefficient = 0.196). With the exception of the Nichols-like clade Laboratory Strain “Chicago” (NC_017268), which was excluded due to a known sequencing error in the reference sequence, all variants were in frame and SNVs limited to the six positions per 60 bp repeat previously recognized to be variable, resulting in four amino acid substitutions. Subclade-specific repeat length variation was noted (*p* < 2×10^−16^, ANOVA), with all but five samples in Nichols subclade B having ten repeats, three of the four strains in Nichols C with 19 repeats, and 15 or 16 repeats in the SS14 Mexico subclade (Figure 2A-B). When samples from Madagascar, which had a bimodal distribution of *arp* repeats, were removed from analysis, no repeat length variation was found among different countries (Figure 2C, *p* > 0.05, ANOVA). Among the 170 samples with 14 ARP repeats, nine different gene sequences, which we have named A-I in order of decreasing prevalence, were represented (Figure 2D), with different usage patterns in different subclades. For example, ARP14 variant A was found in 103 samples in SS14 subclades exclusively. ARP14 variants B and C, which differ by only three nucleotides within a single 60 bp repeat, are found exclusively within the Nichols A subclade, variant E found exclusively in Nichols B, and variant D found in Nichols subclades C, E, and the subclade containing the laboratory strains, Nichols D.

**Figure 2:**
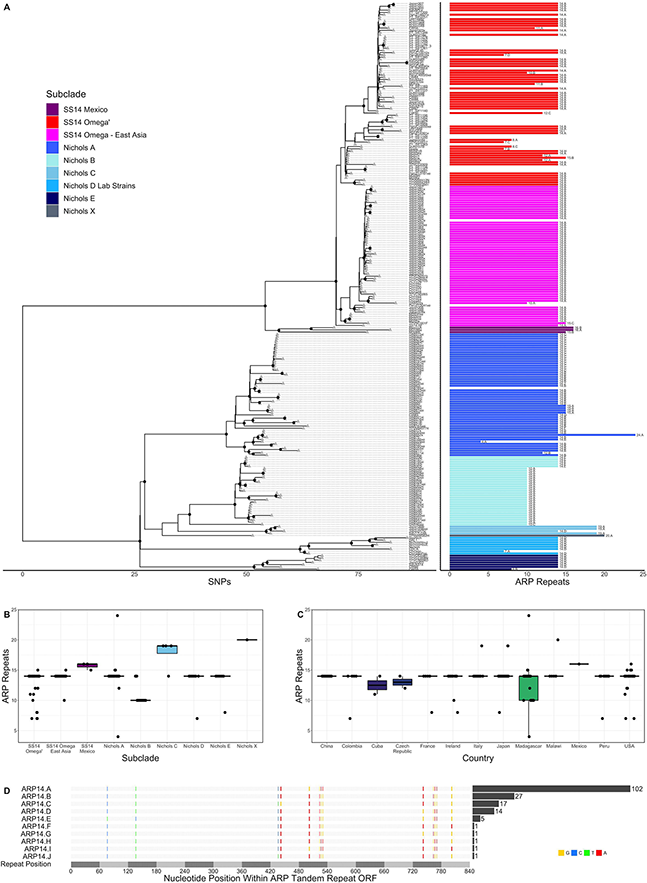
Variation in *arp* repeat length. A) Recombination masked whole genome phylogeny (left) with the number of *arp* repeats for each strain (right). Sequence variant number is included as text to the right of each length bar. All data are also included in a tabular representation in Supplementary Table 2. Number of *arp* repeats by subclade (B) or country (C). D) Multiple sequence alignment of the nine variants with 14 *arp* repeats. Variant positions are highlighted and bases colored red, blue, yellow, or green for A, C, G, or T, respectively. The number of strains with each variant sequence is included in the bar graph to the right of the multiple sequence alignment.

### The sequence of *arp* repeats varies per subclade

We also characterized the pattern of the modular 60 bp near-identical repeats. Three types of *arp* repeat (Type I, II, and III) were originally identified by Liu et al (Liu et al., 2007). A fourth type, Type II/III, which likely formed by recombination of Types II and III between the two sets of three variable positions to form a chimera, was discovered in a larger analysis of laboratory and clinical strains (Harper et al., 2008). In addition to these “canonical” types of *arp* repeat, we found three additional repeat Types that had not been previously described (Fig 3A): Type I/III, which appears to be a chimera of Types I and III and found only in a single Peruvian strain with seven *arp* repeats; Type III/I, which is a chimera of Type III and either Type I or II and found in a 14-repeat *arp* variant found in 17 samples from Madagascar as well as the Cuban strain CW83 (Grillová et al., 2019); and Type IIIG, found in a single sample from the United States, which shares the Type III sequence at the first four variable positions and likely recombination to match Types I or II at the final two variable positions. All repeat Types generated unique amino acid sequences (Fig. 3B). Consistent with previous reports that non-venereal *T. pallidum* subspecies use only Type II repeats (Harper et al., 2008; Cejková et al., 2012; Staudová et al., 2014), the single Lihir Island *T. pallidum* subsp. *pertenue* strain and four Japan *T. pallidum* subsp. *endemicum* strains included in our previous study (Lieberman et al., 2021) had only four Type II repeats, or ten or eleven Type II repeats, respectively (Supporting Information).

**Figure 3:**
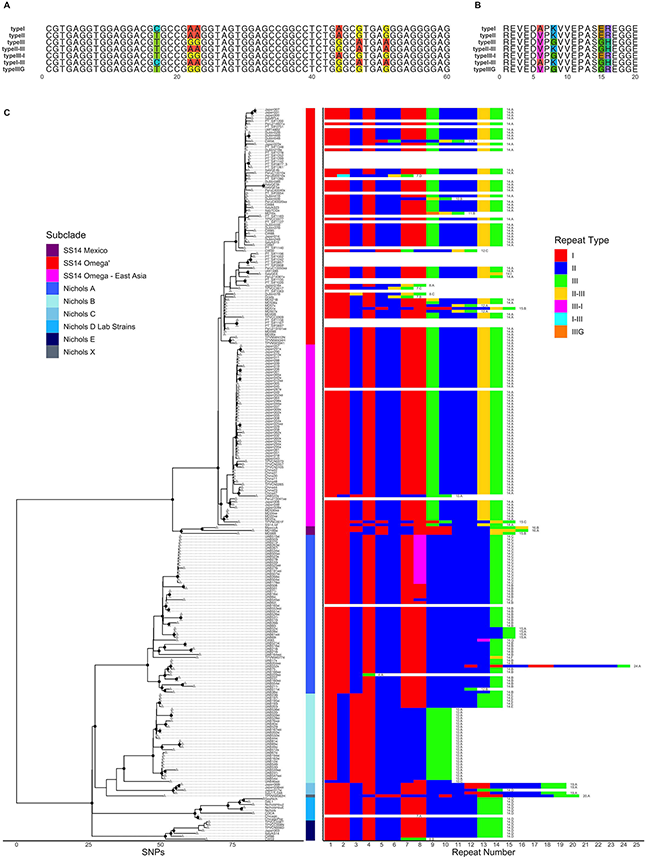
*arp* repeat type usage. A) Nucleotide sequence of the arp repeat module types. Variable positions are highlighted. B) Amino acid sequence of the arp repeat module types. Variable positions are highlighted. C) Recombination masked whole genome phylogeny (left) with the repeat type usage per strain (right). The 60 bp arp repeats are colored by type.

To better visualize the relationships and pattern of repeat Type use between different *arp* variants, we plotted them in the context of the whole genome phylogeny (Fig. 3C). In this context, several general patterns emerge. Most strikingly, out of 121 strains in the SS14-like clade with ARP sequence information, 115 include least two Type II repeats, a penultimate Type II/III repeat, and a 3’ Type III repeat, in contrast to the Nichols-like clade, which instead contains four or more Type II repeats followed by one (Nichols subclade A) or two (Nichols subclades B-E) Type III repeats. Furthermore, with the exception of *arp* sequences in the Nichols B subclade, which start with a Type I repeat followed by Type II, *arp* variants in 99% of strains across both major clades start with two Type I repeats.

Within each subclade, we defined the dominant *arp* sequence as the one most commonly found. Out of the 226 *T. pallidum* subsp. *pallidum* strains with *arp* sequence information, we identified 170 strains with the dominant *arp* sequence in the subclade to which it belongs, 25 strains where the *arp* repeat sequence clearly did not match the dominant sequence in its subclade, and 26 strains that comprised three clusters of closely-related strains all exhibiting the same *arp* variant but diverged from the predominant variant in the subclade to which they belonged. (“Dominant *arp* Variant in Subclade”, Supporting Information). The dominant strain in the SS14 Mexico or Nichols X subclades could not be determined, and Nichols D Lab Strain Chicago was excluded due to a known sequencing error. We did not find any SNVs enriched among strains with an *arp* sequence altered from the dominant strain (p>0.05, Fisher’s exact test). We also examined the relationship between non-dominant *arp* sequence and branch length on the whole genome maximum likelihood tree. We found that the strains with the non-dominant *arp* sequence had terminal branch lengths (defined as the number of SNPs that separate a tip from its most recent ancestral node) that were on average 3.6 times longer than those with the dominant sequence (SFig4, mean (median) 4.11 (3) SNPs vs 1.14 (0) SNPs, *p*=0.0046, Welch’s t-test). While these observations do not account for sources of selective pressure such as host immune response or anatomic site of infection, they do suggest that events that result in a novel *arp* sequence are likely stochastic and less frequent than SNP fixation, which occurs approximately once per genome every five years (Beale et al., 2019; Lieberman et al., 2021; Taouk et al., 2022).

### Novel *arp* sequences likely arise through intra- or inter-strain recombination

In cases where a strain’s *arp* sequence did not match the dominant sequence in its subclade, we confirmed the sequence by examination of WGS short read linkage and/or Sanger sequencing. We then attempted to determine the simplest mechanism to generate the novel sequence. In most cases, a single intra-strain recombination event, resulting in insertion, deletion, or substitution of one or more *arp* modules, is most parsimonious (SFig5; most variants could theoretically be generated by addition or removal of modules from the dominant sequence in a different pattern than shown). For example, in the SS14 Omega’
s and East Asia subclades, ARP14.A is most commonly modified presumptively through loss of repeat modules, resulting in sequence lengths between 7 and 12 modules. Notably, though, we do not know how long the recombination junctions must be, therefore sequences such as ARP11.B, which could have been formed by a recombination event between the 4^th^ and 5^th^ variable nucleotides of type III and type II modules as shown, or could also have been generated via deletion of the three tandem type II repeats in conjunction with mutation of the 5^th^ and 6^th^ variable positions in the type III repeat from A to G, to generate the novel Type IIIG module.

In addition to novel *arp* variants that can be generated via a single intrastrain recombination, there are other variants whose presumptive lineage is less clear. For example, the strain UAB46xei is very unusual, both starting and ending with type II modules, unlike any other *arp* variants. Interestingly, though, the UAB46xei *arp* sequence is 10 repeats, like the ARP10.A sequence that predominates the Nichols B subclade. Strains TPVMW082H and Dublin57B, belonging to Nichols X and SS14 Omega’
s subclades, respectively, also contain unique sequences generated via complex mechanisms, though it is plausible that several individual recombination events generated the 20-repeat variant found in TPVMW082H, which is only distantly related to any other strains in this dataset.

Other strains show clear evidence of inter-strain recombination. For example, ARP10.A is the dominant sequence in the Madagascar strains that comprise Nichols subclade B and found in no other strains except for the single Madagascar sample in the SS14 East Asia subclade. Similarly, although the dominant *arp* variant in the SS14 Mexico subclade cannot be determined since each strain has a unique sequence, the *arp* variant in SS14 Mexico strain MD06B is only shared with the SS14 Omega’
s subclade strain MD51x, both of which were collected from Maryland, USA. Finally, strain Japan317x in Nichols subclade C harbors the same ARP14.D variant as is found in strains in Nichols subclades D and E, including in one Japan sample; however, deletion of 5 modules from the ARP19.A variant private to the Nichols C subclade is also a possible mechanism for generation of the 14.D variant in Japan317x. Together, these data suggest that both intra- and inter-strain recombination is employed by *T. pallidum* to generate diversity at the *arp* locus.

We also attempted to determine if *tp0470* repeat length or *arp* repeat length and sequence were associated with syphilis stage. Among the 79 strains with stage information available, 49 were primary and 30 were secondary; although longer *tp0470* variants were seen on average in secondary syphilis samples (SFig6A, mean (median) 23.6 (25) for secondary vs 18.8 (15) for primary, *p*=0.03624, Welch’s t-test), no significant differences in *tp0470* repeat length by disease stage were observed when samples were further split by SS14- or Nichols-like clades, suggesting that sampling bias may be confounding interpretation of the association of *tp0470* repeat length with disease stage. There was no association between non-dominant *arp* sequence and primary or secondary syphilis (p>0.05, Fisher’s exact test), nor did we find a significant difference in the number of *arp* repeats among primary vs secondary syphilis samples (Sfig6B, *p*=0.358, Welch’s t-test). However, secondary syphilis was overrepresented among strains with the ARP10.A sequence (Fisher’s Exact Test, *p*=0.0171), while primary syphilis was overrepresented among strains with the ARP14.A sequence (Fisher’s Exact Test, *p*=0.0188). No other *arp* sequence variants had enough samples with stage data to determine overrepresentation.

### Structural modeling of ARP and TP0470 to localize 3D repeat structure

There is ample evidence the proteins encoded by *tp0470* and *arp* genes are present during infection: Previous studies have shown that the *tp0470* transcript is expressed (Smajs et al., 2005; De Lay et al., 2021), and sera from infected rabbits (McKevitt et al., 2005) and patients (Brinkman et al., 2006) are reactive to TP0470 protein. The *arp* transcript is expressed (Smajs et al., 2005; De Lay et al., 2021), and the ARP protein was found to be one of the top 10% most abundant proteins by mass spectrometry (Osbak et al., 2016). *T. pallidum*-infected rabbit sera are reactive to ARP (McKevitt et al., 2005; Liu et al., 2007), while sera from infected human patients are weakly reactive to ARP during primary infection (Brinkman et al., 2006). Therefore, we attemped to model select variants of the full-length proteins encoded by *tp0470* and *arp*. The *tp0470* gene is identical in all strains included in this study outside of the repeat length variation, and TP0470 is confidently predicted by both SignalP 6.0 (Teufel et al., 2022) and PSORTb V3.0 (Yu et al., 2010) to contain a signal sequence with no lipid anchor, which suggests it resides in the periplasm. A conserved domain search (Lu et al., 2020) reveals the tetratricopeptide repeat protein domain (e-value: 1.52e^-8^) at the N-term of the protein, with no predicted conserved domains otherwise. This is consistent in structures predicted by both trRosetta (Yang et al., 2020; Du et al., 2021) and AlphaFold (Jumper et al., 2021; Varadi et al., 2022), which contain four pairs of antiparallel α-helices that comprise the conserved tetratricopeptide motif in the N terminus, followed by an extended α-helix largely composed of the highly charged eight amino acid repeat motif “EAEEARRK” (Fig 4A-B; tetratricopeptide repeat motifs in green, pre-repeat linker in grey, 15 repeats of eight amino acids in purple, post-repeat C terminus in gold). Confidence metrics for trRosetta are high for the overall structure (TM-score = 0.756), while for AlphaFold the local Difference Distance Test score is >80 (high) throughout the tetratricopeptide repeat domain, and drops throughout the length of the extended helix. Variants with longer tandem repeats are predicted to have a helix that folds back on itself by AlphaFold, while trRosetta predicts an elongated structure; although at low confidence (SFig7A-B). The length of the repeat portion of the helix ranges between 4-51 repeats, or a total of 32-408 residues in repeats, with a modal number of repeats of 15 (SFig7C). Assuming 0.54nm in length per helical turn of 3.6 residues, the elongated length of the predicted helix may range between 11.5 nm and 68.1 nm including the non-repetitive 23 amino acids N terminal and 23 amino acids C terminal to the repeats, with 90% of lengths between 7-36 repeats (15.3-50.1 nm), and the helix of the 15 repeat variant measuring approximately 24.9 nm. Within the helix, a single eight-residue repeat makes just over two helical turns. Fig 4C zooms in on four repeats, comprising approximately 9 helical turns. Although the orientation of amino acid residues in a structural model does not reflect the precise native conformation, the stick representation of sidechains (Fig 4C, top) and smoothed surface charge (Fig 4C, bottom) demonstrates the highly polar nature of the TP0470 repeats.

**Figure 4:**
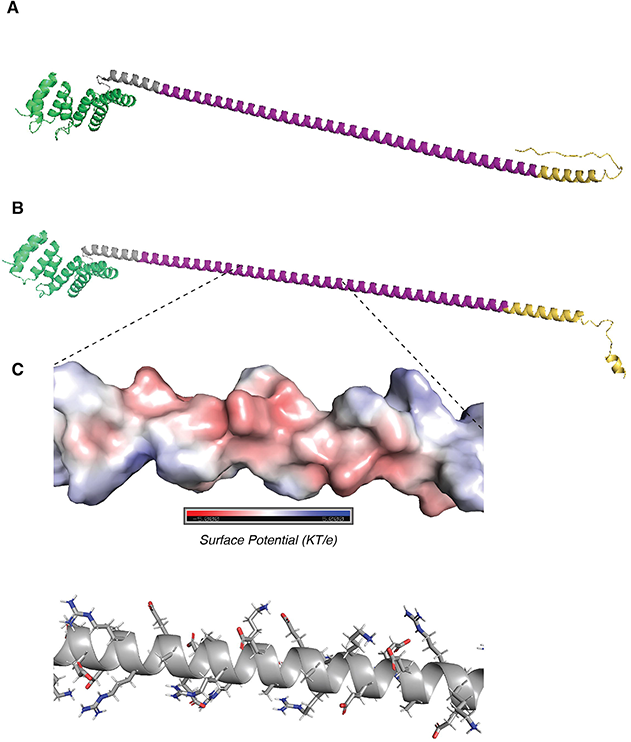
Structure predictions of TP0470. A) trRosetta and B) AlphaFold predictions of structure of 15 repeat TP0470 variant. N terminal tetratricopeptide repeat domain is shown in green, repeats are in purple, and C-terminal region is in gold. C) APBS electrostatic surface potential (top) and stick model of sidechains (bottom) for portion of repeat helix.

We predicted domains and structures for select variants of the acidic repeat protein. By SignalP 6.0, it is predicted to have a signal sequence (probability=0.67) and possibly lipidation site at cysteine-29 (probability=0.33), however, PSORT predicts neither of these elements. A conserved domain search reveals a C-terminal SPOR domain (e value: 7.5e^-3^), which in other proteins is a peptidoglycan binding domain (Yahashiri et al., 2017). Together, these results suggest that the acidic repeat protein is localized to the periplasm.

For model generation, we first examined the ARP14.A variant, by far the most common variant in our phylogeny, harbored by 103 strains. While both trRosetta and AlphaFold predicted the expected twisted beta strand structure of the C-terminal SPOR domain (Fig 5A-B, magenta), AlphaFold’s low confidence prediction of the repeat regions (local Distance Difference Test ∼40) is entirely unstructured (Fig 5B), whereas the trRosetta prediction is for the acidic repeats to form a disordered linker comprising the first five repeats, followed by a parallel β-sheet structure that contains nine strands composed of the last nine acidic repeats (Fig 5A), although the confidence in the prediction is quite low (TM-score=0.288). This structure would contain an extremely acidic face of the β-sheet (Fig 5C) with a periodicity of 20 residues, the same as the repeat. Interestingly, the repeat region in most other variants was not predicted by trRosetta to fold into a beta sheet, rather, they were highly disordered (SFig8); only variants ARP14.H and ARP15.A were also predicted to form a β-sheet from the repeats. Overall, despite a plausible structure for some variants, structural modeling of the acidic repeat protein remains challenging with only low confidence models returned by two methods and any attempt to infer function based on these results should be made cautiously.

**Figure 5:**
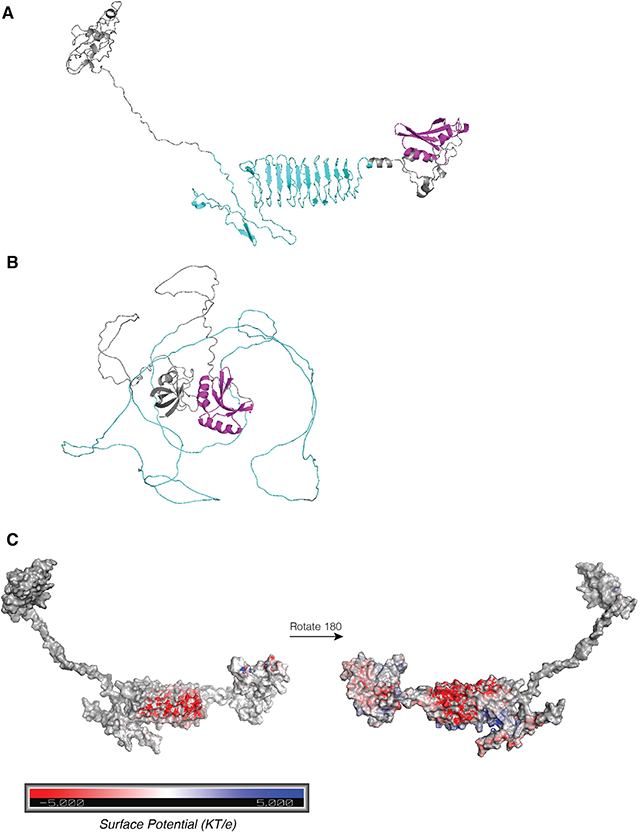
Structure predictions ARP14A. A) trRosetta and B) AlphaFold predictions of structure of ARP14A. C-terminal SPOR domain is shown in magenta, repeats in cyan. C) APBS electrostatic surface potential for trRosetta ARP structure from A. Red denotes negative charge (acidic) and blue denotes positive charge (basic).

## Discussion

Despite the relatively low rate of SNP fixation, with a mean rate of approximately 1-3×10^−7^ substitutions per site per year in putative non-recombinogenic loci (Beale et al., 2019; Lieberman et al., 2021; Taouk et al., 2022)), *T. pallidum* uses additional mechanisms to increase its genetic diversity and antigenic repertoire. These include inter-strain and inter-species recombination in genes encoding the Tpr family of antigens (Gray et al., 2006; Kumar et al., 2018; Grillová et al., 2019), gene conversion in the variable regions of *tprK* (Centurion-Lara et al., 2004; Giacani et al., 2010; Reid et al., 2014), and homopolymer expansion and contraction to alter promoter activity and hence expression level of putative outer membrane proteins (Giacani et al., 2015). Previous work has demonstrated that diversity in the *tp0470* and *arp* repeat length, and repeat type usage in *arp*, is likely generated by recombination, although modification of the number of repeats in the *tp0470* could also be possible via a polymerase slippage mechanism. Our current study extends these findings to a large cohort of clinical samples with near-complete genomes available, enabling examination of differences between subclades and correlation with genome features.

From our results, it is clear that a very wide distribution of *tp0470* repeat lengths is possible but with no sequence variation within the repeat. Although *T. pallidum* has an extremely low rate of mutation and most genes are highly conserved, the absence of sequence variation within the *tp0470* gene outside of repeat length variation suggests the protein may be under purifying selection. The *arp* gene has multiple sequence variants generated by using different repeat module types in a tandem arrangement, but is highly enriched for sequences with fourteen repeats. For both genes, some differences between Nichols- and SS14-like clades are observed: in *tp0470*, there is a slight increase in repeat length in the Nichols-like clade vs SS14. In the *arp* gene, variants are generally limited to a single subclade, particularly in the Nichols-like clade, which has far greater genetic diversity than the SS14-like clade, with an average pairwise SNP distance of 42, as compared to an average pairwise SNP distance of 10 for the SS14-like clade. It is unclear whether differences between *tp0470* lengths or repeat module pattern in *arp* between subclades have functional consequences and are being selected for, or whether the differences simply reflect random events during diversification. In the case of *arp*, we did not find any SNPs throughout the genome that correlated with an unexpected repeat sequence.

Until very recently (Romeis et al., 2021), no reverse genetics system for *T. pallidum* existed, therefore, traditional bacteriological genetic tools, such as mutants and knockout strains, to interrogate gene functions have not been available for the syphilis spirochete. Prior to the development of an epithelial cell co-culture system in 2018 (Edmondson et al., 2018), *T. pallidum* could only be passaged through rabbit testes, precluding forward genetics screens. While proteome-wide bioinformatic structural predictions have helped to shed light on the likely role of conserved structural domains (Houston et al., 2018), the structure and function *T. pallidum* proteins containing novel motifs, such as the repeat sequences found in TP0470 and ARP, remain unknown. To gain insight into their biological function and possible role in syphilis pathogenesis, we employed several *in silico* tools to predict the topology and structure of full-length ARP and TP0470 proteins.

The presence of a signal peptide on TP0470 is strong evidence that it is localized to the periplasm, where it likely binds other as-yet-undermined protein(s) via its N-terminal tetratricopeptide motif, which was predicted by both trRosetta and AlphaFold. Both algorithms predicted an extended α-helix with regions of alternating positive and negative surface electrostatic potential, regardless of the length of the repeats. Although TP0470 does not have any known interacting partners, it seems plausible that in addition to interactions formed by tetratricopeptide motif, the unusually long, very polar α-helix that comprises the repeats also serves to mediate protein-protein interactions.

Although a signal peptide was not confidently predicted by one tool (SignalP 6.0) and not predicted at all by a second (PSORT), the presence of a C-terminal SPOR domain, which binds denuded peptidoglycan, strongly suggests ARP must be present in the periplasm. However, it is less clear whether ARP is free in the periplasm or is acylated at cys-29 (weakly predicted by SignalP 6.0), tethering it to the inner membrane. Both topologies are consistent with other SPOR domain-containing proteins (Yahashiri et al., 2017); mass spectrometry, Edman degradation, or other biochemical techniques will be necessary to resolve this question.

In addition to the unclear localization of ARP, the structure formed by the ARP repeat modules remains murky. The most biologically plausible structure generated by the modeling software is of the modular repeats forming a parallel beta sheet, with periodicity of 20 residues, the same length as the repeats. The β-sheet and the loops that connect the strands form an extremely negatively charged surface; it seems likely that whatever the ARP repeat domain binds, it will be positively charged.

Figure 6 summarizes our current understanding of the structure and topology of TP0470 and ARP. Because TP0470 is predicted to be soluble, it may be able to traverse the peptidoglycan layer through pores. In contrast, ARP may be associated with the inner membrane, constraining its movement within the periplasm. However, these models do not resolve why there is tremendous diversity of *tp0470* repeat lengths, and *arp* repeat lengths and repeat module usage. To answer these questions and determine the biological function of the repeat domains, extensive biochemical and biophysical studies of different variants will be necessary.

**Figure 6:**
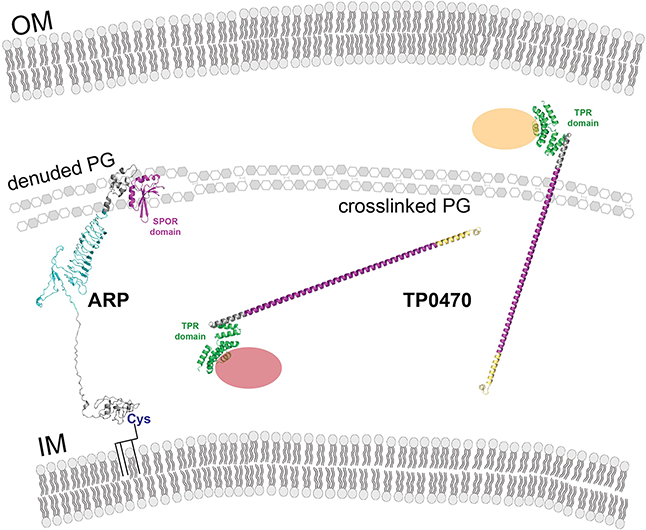
Model showing ARP and TP0470 cellular location and putative interactions. Both ARP and TP0470 are localized to the periplasm. The ARP N terminus may be acylated at cysteine 29. OM: Outer Membrane. IM: Inner Membrane. PG: Peptidoglycan.

One of the primary limitations of our genomic dataset is that of bias introduced by unequal sampling. For example, although we and others have reported that 14-repeat arp variants predominate, particularly variant 14.A, this may reflect the increased sampling in geographical regions (Europe and the United States) where SS14-like clade strains predominate. Furthermore, many of the strains included in our study were not collected with extensive clinical histories or patient characteristics, limiting our ability to infer functional differences from sequence variation. Finally, the use of PCR to interrogate genes containing repetitive sequences is always challenging due to the generation of truncated artifact products, which can amplify preferentially over the “real” product, as we saw for *tp0470* in some samples (SFig1B). However, these products were readily apparent in both the gel images and histograms of mapped reads; therefore the data remained interpretable, and were consistent in technical replicates (Supporting information).

In addition to developing a novel method to examine repeat length and sequence in two challenging genomic loci in *T. pallidum*, our study has demonstrated extensive *tp0470* and *arp* repeat diversity among more than 200 clinical strains with whole genome sequence, by far the largest study of these genes to date. Importantly, we found that more than 10% of strains contained an *arp* variant that had a different length and sequence than the dominant variant in the subclade, which builds on concerns about the utility of using the number of *arp* repeats as part of strain typing tools for epidemiology (Mikalová et al., 2013). Finally, we have proposed a possible mechanism by which each may interact with peptidoglycan and/or other periplasmic factors and influence morphogenesis. Although additional genetic, biophysical, and biochemical interaction studies will be necessary to characterize their function and elucidate their binding partners, our study of *tp0470* and *arp* lays the foundation to directly link the genotype to function of two novel genes that may influence *T. pallidum* pathogenesis.

## Data Availability

Sequencing data is available under NCBI BioProjects PRJNA723099 and PRJNA815321. Supporting Information contains BioSample and accession information for nanopore reads.

## Funding

This work was supported, in whole or in part, by the Bill & Melinda Gates Foundation INV-036560. Under the grant conditions of the Foundation, a Creative Commons Attribution 4.0 Generic License has already been assigned to the Author Accepted Manuscript version that might arise from this submission (ALG, LG, JBP, ACS, JDR, and KLH). This work was further supported by NIAID grant numbers U19AI144133 (LG and ALG) and U19AI144177 (JDR, KLH, JCS, ACS, and JBP), as well as grants from Ministry of Education, Culture, Sports, Science and Technology of Japan (number 21K09388) to SN and from Japan Agency for Medical Research and Development (number 21fk0108091j0303) to MO. Additional research funds were generously provided by Connecticut Children’s (JCS, JDR, and KLH).

## Figure Legends

**Figure S1: *tp0470* PCR band and nanopore pipeline call concordance of select samples**. A) PCR bands following barcoding. Bands should have a size of 24 bp times the number of repeats plus 273, including both flanking regions (225 bp) and two barcodes (48 bp total). The red box highlights non-specific amplification of low molecular weight fragments, while the red arrow shows the correct band size. B) Histogram of the distribution of reads aligned to each length variant in the mapping reference file.

**Figure S2: *arp* PCR band and nanopore pipeline call concordance of select samples**. A) PCR bands following barcoding. Bands should have a size of 60 bp times the number of repeats plus 874, including both flanking regions (826 bp) and two barcodes (48 bp total). B) Histogram of the distribution of reads aligned to each length variant in the mapping reference file.

**Figure S3: *arp* repeat length is not correlated with *tp0470* repeats**. No correlation was seen between number of *arp* and *tp0470* repeats (Pearson coefficient = 0.008).

**Figure S4: Terminal branch lengths are longer for strains with non-dominant arp sequences**. Average branch lengths from tip to ancestral node were determined for strains with dominant and non-dominant arp variants. **, *p*=0.0046, Welch’s t-test.

**Figure S5: Possible single recombination events to generate arp variants from the dominant arp variant in each subclade**. The dominant sequence from each subclade is shown in the top position for each pair, and the non-dominant (possibly nascent) variant below. Dotted lines show possible junctions. Only a single possibility per variant is shown.

**Figure S6: Longer *tp0470* repeats are associated with secondary syphilis**. Out of 79 samples with stage information, samples from secondary syphilis had on average approximately 5 more repeats than primary (*, *p*=0.03624, Welch’s t-test).

**Figure S7: TP0470 predictions**: A) trRosetta and B) AlphaFold predictions of structure of 51 repeat TP0470 variant. Structures are colored from blue to red N term to C term. C) Distribution of *tp0470* variants in phylogeny.

**Figure S8: trRosetta predictions for additional ARP variants**. Structures are colored from blue to red N term to C term.

## Notes

### Competing Interest Statement

The authors have declared no competing interest.

## References

Arora, N., Schuenemann, V. J., Jäger, G., Peltzer, A., Seitz, A., Herbig, A., et al. (2017). Origin of modern syphilis and emergence of a pandemic Treponema pallidum cluster. Nat Microbiol 2, 16245. doi: 10.1038/nmicrobiol.2016.245.

Beale, M. A., Marks, M., Cole, M. J., Lee, M.-K., Pitt, R., Ruis, C., et al. (2021). Global phylogeny of Treponema pallidum lineages reveals recent expansion and spread of contemporary syphilis. Nat Microbiol 6, 1549–1560. doi: 10.1038/s41564-021-01000-z.

Beale, M. A., Marks, M., Sahi, S. K., Tantalo, L. C., Nori, A. V., French, P., et al. (2019). Genomic epidemiology of syphilis reveals independent emergence of macrolide resistance across multiple circulating lineages. Nat Commun 10, 3255. doi: 10.1038/s41467-019-11216-7.

Brinkman, M. B., McKevitt, M., McLoughlin, M., Perez, C., Howell, J., Weinstock, G. M., et al. (2006). Reactivity of antibodies from syphilis patients to a protein array representing the Treponema pallidum proteome. J Clin Microbiol 44, 888–891. doi: 10.1128/JCM.44.3.888-891.2006.

Bushnell, B. BBMap short read aligner, and other bioinformatic tools.

Cejková, D., Zobaníková, M., Chen, L., Pospíšilová, P., Strouhal, M., Qin, X., et al. (2012). Whole genome sequences of three Treponema pallidum ssp. pertenue strains: yaws and syphilis treponemes differ in less than 0.2% of the genome sequence. PLoS Negl Trop Dis 6, e1471. doi: 10.1371/journal.pntd.0001471.

Centurion-Lara, A., LaFond, R. E., Hevner, K., Godornes, C., Molini, B. J., Van Voorhis, W. C., et al. (2004). Gene conversion: a mechanism for generation of heterogeneity in the tprK gene of Treponema pallidum during infection. Mol Microbiol 52, 1579–1596. doi: 10.1111/j.1365-2958.2004.04086.x.

Cerveny, L., Straskova, A., Dankova, V., Hartlova, A., Ceckova, M., Staud, F., et al. (2013). Tetratricopeptide repeat motifs in the world of bacterial pathogens: role in virulence mechanisms. Infect Immun 81, 629–635. doi: 10.1128/IAI.01035-12.

Chen, W., Šmajs, D., Hu, Y., Ke, W., Pospíšilová, P., Hawley, K. L., et al. (2021). Analysis of Treponema pallidum Strains From China Using Improved Methods for Whole-Genome Sequencing From Primary Syphilis Chancres. J Infect Dis 223, 848–853. doi: 10.1093/infdis/jiaa449.

Currin, A., Swainston, N., Dunstan, M. S., Jervis, A. J., Mulherin, P., Robinson, C. J., et al. (2019). Highly multiplexed, fast and accurate nanopore sequencing for verification of synthetic DNA constructs and sequence libraries. Synthetic Biology 4, ysz025. doi: 10.1093/synbio/ysz025.

De Lay, B. D., Cameron, T. A., De Lay, N. R., Norris, S. J., and Edmondson, D. G. (2021). Comparison of transcriptional profiles of Treponema pallidum during experimental infection of rabbits and in vitro culture: Highly similar, yet different. PLoS Pathog 17, e1009949. doi: 10.1371/journal.ppat.1009949.

Du, Z., Su, H., Wang, W., Ye, L., Wei, H., Peng, Z., et al. (2021). The trRosetta server for fast and accurate protein structure prediction. Nat Protoc 16, 5634–5651. doi: 10.1038/s41596-021-00628-9.

Edmondson, D. G., Hu, B., and Norris, S. J. (2018). Long-Term In Vitro Culture of the Syphilis Spirochete Treponema pallidum subsp. pallidum. mBio 9. doi: 10.1128/mBio.01153-18.

Flasarová, M., Pospíšilová, P., Mikalová, L., Vališová, Z., Dastychová, E., Strnadel, R., et al. (2012). Sequencing-based molecular typing of treponema pallidum strains in the Czech Republic: all identified genotypes are related to the sequence of the SS14 strain. Acta Derm Venereol 92, 669–674. doi: 10.2340/00015555-1335.

Giacani, L., Brandt, S. L., Ke, W., Reid, T. B., Molini, B. J., Iverson-Cabral, S., et al. (2015). Transcription of TP0126, Treponema pallidum putative OmpW homolog, is regulated by the length of a homopolymeric guanosine repeat. Infect Immun 83, 2275–2289. doi: 10.1128/IAI.00360-15.

Giacani, L., Molini, B. J., Kim, E. Y., Godornes, B. C., Leader, B. T., Tantalo, L. C., et al. (2010). Antigenic variation in Treponema pallidum: TprK sequence diversity accumulates in response to immune pressure during experimental syphilis. J Immunol 184, 3822–3829. doi: 10.4049/jimmunol.0902788.

Gray, R. R., Mulligan, C. J., Molini, B. J., Sun, E. S., Giacani, L., Godornes, C., et al. (2006). Molecular evolution of the tprC, D, I, K, G, and J genes in the pathogenic genus Treponema. Mol Biol Evol 23, 2220–2233. doi: 10.1093/molbev/msl092.

Grillová, L., Oppelt, J., Mikalová, L., Nováková, M., Giacani, L., Niesnerová, A., et al. (2019). Directly Sequenced Genomes of Contemporary Strains of Syphilis Reveal Recombination-Driven Diversity in Genes Encoding Predicted Surface-Exposed Antigens. Front Microbiol 10, 1691. doi: 10.3389/fmicb.2019.01691.

Grimes, M., Sahi, S. K., Godornes, B. C., Tantalo, L. C., Roberts, N., Bostick, D., et al. (2012). Two mutations associated with macrolide resistance in Treponema pallidum: increasing prevalence and correlation with molecular strain type in Seattle, Washington. Sex Transm Dis 39, 954–958. doi: 10.1097/OLQ.0b013e31826ae7a8.

Harper, K. N., Liu, H., Ocampo, P. S., Steiner, B. M., Martin, A., Levert, K., et al. (2008). The sequence of the acidic repeat protein (arp) gene differentiates venereal from nonvenereal Treponema pallidum subspecies, and the gene has evolved under strong positive selection in the subspecies that causes syphilis. FEMS Immunol Med Microbiol 53, 322–332. doi: 10.1111/j.1574-695X.2008.00427.x.

Hook III, E. W., Behets, F., Van Damme, K., Ravelomanana, N., Leone, P., Sena, A. C., et al. (2010). A Phase III Equivalence Trial of Azithromycin versus Benzathine Penicillin for Treatment of Early Syphilis. J INFECT DIS 201, 1729–1735. doi: 10.1086/652239.

Hopkins, S., Lyons, F., Coleman, C., Courtney, G., Bergin, C., and Mulcahy, F. (2004). Resurgence in Infectious Syphilis in Ireland: An Epidemiological Study. Sexually Transmitted Diseases 31, 317–321. doi: 10.1097/01.OLQ.0000123653.84940.59.

Houston, S., Lithgow, K. V., Osbak, K. K., Kenyon, C. R., and Cameron, C. E. (2018). Functional insights from proteome-wide structural modeling of Treponema pallidum subspecies pallidum, the causative agent of syphilis. BMC Struct Biol 18, 7. doi: 10.1186/s12900-018-0086-3.

Jumper, J., Evans, R., Pritzel, A., Green, T., Figurnov, M., Ronneberger, O., et al. (2021). Highly accurate protein structure prediction with AlphaFold. Nature 596, 583–589. doi: 10.1038/s41586-021-03819-2.

Kumar, S., Caimano, M. J., Anand, A., Dey, A., Hawley, K. L., LeDoyt, M. E., et al. (2018). Sequence Variation of Rare Outer Membrane Protein β-Barrel Domains in Clinical Strains Provides Insights into the Evolution of Treponema pallidum subsp. pallidum, the Syphilis Spirochete. mBio 9. doi: 10.1128/mBio.01006-18.

Li, H., and Durbin, R. (2009). Fast and accurate short read alignment with Burrows-Wheeler transform. Bioinformatics 25, 1754–1760. doi: 10.1093/bioinformatics/btp324.

Lieberman, N. A. P., Lin, M. J., Xie, H., Shrestha, L., Nguyen, T., Huang, M.-L., et al. (2021). Treponema pallidum genome sequencing from six continents reveals variability in vaccine candidate genes and dominance of Nichols clade strains in Madagascar. PLoS Negl Trop Dis 15, e0010063. doi: 10.1371/journal.pntd.0010063.

Liu, D., He, S.-M., Zhu, X.-Z., Liu, L.-L., Lin, L.-R., Niu, J.-J., et al. (2020). Molecular Characterization Based on MLST and ECDC Typing Schemes and Antibiotic Resistance Analyses of Treponema pallidum subsp. pallidum in Xiamen, China. Front Cell Infect Microbiol 10,618747. doi: 10.3389/fcimb.2020.618747.

Liu, H., Rodes, B., George, R., and Steiner, B. (2007). Molecular characterization and analysis of a gene encoding the acidic repeat protein (Arp) of Treponema pallidum. J Med Microbiol 56, 715–721. doi: 10.1099/jmm.0.46943-0.

Lu, S., Wang, J., Chitsaz, F., Derbyshire, M. K., Geer, R. C., Gonzales, N. R., et al. (2020). CDD/SPARCLE: the conserved domain database in 2020. Nucleic Acids Res 48, D265–D268. doi: 10.1093/nar/gkz991.

Lukehart, S. A., Godornes, C., Molini, B. J., Sonnett, P., Hopkins, S., Mulcahy, F., et al. (2004). Macrolide resistance in Treponema pallidum in the United States and Ireland. N Engl J Med 351, 154–158. doi: 10.1056/NEJMoa040216.

Marra, C. M., Sahi, S. K., Tantalo, L. C., Godornes, C., Reid, T., Behets, F., et al. (2010). Enhanced molecular typing of treponema pallidum: geographical distribution of strain types and association with neurosyphilis. J Infect Dis 202, 1380–1388. doi: 10.1086/656533.

McKevitt, M., Brinkman, M. B., McLoughlin, M., Perez, C., Howell, J. K., Weinstock, G. M., et al. (2005). Genome scale identification of Treponema pallidum antigens. Infect Immun 73, 4445–4450. doi: 10.1128/IAI.73.7.4445-4450.2005.

Mikalová, L., Pospíšilová, P., Woznicová, V., Kuklová, I., Zákoucká, H., and Smajs, D. (2013). Comparison of CDC and sequence-based molecular typing of syphilis treponemes: tpr and arp loci are variable in multiple samples from the same patient. BMC Microbiol 13, 178. doi: 10.1186/1471-2180-13-178.

Molepo, J., Pillay, A., Weber, B., Morse, S. A., and Hoosen, A. A. (2007). Molecular typing of Treponema pallidum strains from patients with neurosyphilis in Pretoria, South Africa. Sex Transm Infect 83, 189–192. doi: 10.1136/sti.2006.023895.

Noda, A. A., Méndez, M., Rodríguez, I., and Šmajs, D. (2022). Genetic Recombination in Treponema pallidum: Implications for Diagnosis, Epidemiology, and Vaccine Development. Sex Transm Dis 49, e7–e10. doi: 10.1097/OLQ.0000000000001497.

Osbak, K. K., Houston, S., Lithgow, K. V., Meehan, C. J., Strouhal, M., Šmajs, D., et al. (2016). Characterizing the Syphilis-Causing Treponema pallidum ssp. pallidum Proteome Using Complementary Mass Spectrometry. PLoS Negl Trop Dis 10, e0004988. doi: 10.1371/journal.pntd.0004988.

Parr, J. B. (2021). IDEELResearch/tpallidum_genomics: Tpallidum_genomics. doi: 10.5281/ZENODO.5773174.

Pětrošová, H., Zobaníková, M., Čejková, D., Mikalová, L., Pospíšilová, P., Strouhal, M., et al. (2012). Whole genome sequence of Treponema pallidum ssp. pallidum, strain Mexico A, suggests recombination between yaws and syphilis strains. PLoS Negl Trop Dis 6, e1832. doi: 10.1371/journal.pntd.0001832.

Pillay, A., Liu, H., Chen, C. Y., Holloway, B., Sturm, A. W., Steiner, B., et al. (1998). Molecular subtyping of Treponema pallidum subspecies pallidum. Sex Transm Dis 25, 408–414. doi: 10.1097/00007435-199809000-00004.

Pillay, A., Liu, H., Ebrahim, S., Chen, C. Y., Lai, W., Fehler, G., et al. (2002). Molecular typing of Treponema pallidum in South Africa: cross-sectional studies. J Clin Microbiol 40, 256–258. doi: 10.1128/JCM.40.1.256-258.2002.

Pinto, M., Borges, V., Antelo, M., Pinheiro, M., Nunes, A., Azevedo, J., et al. (2016). Genome-scale analysis of the non-cultivable Treponema pallidum reveals extensive within-patient genetic variation. Nat Microbiol 2, 16190. doi: 10.1038/nmicrobiol.2016.190.

Pope, V., Fox, K., Liu, H., Marfin, A. A., Leone, P., Seña, A. C., et al. (2005). Molecular subtyping of Treponema pallidum from North and South Carolina. J Clin Microbiol 43, 3743–3746. doi: 10.1128/JCM.43.8.3743-3746.2005.

Reid, T. B., Molini, B. J., Fernandez, M. C., and Lukehart, S. A. (2014). Antigenic variation of TprK facilitates development of secondary syphilis. Infect Immun 82, 4959–4967. doi: 10.1128/IAI.02236-14.

Romeis, E., Tantalo, L., Lieberman, N., Phung, Q., Greninger, A., and Giacani, L. (2021). Genetic engineering of Treponema pallidum subsp. pallidum, the Syphilis Spirochete. PLoS Pathog 17, e1009612. doi: 10.1371/journal.ppat.1009612.

Smajs, D., McKevitt, M., Howell, J. K., Norris, S. J., Cai, W.-W., Palzkill, T., et al. (2005). Transcriptome of Treponema pallidum: gene expression profile during experimental rabbit infection. J Bacteriol 187, 1866–1874. doi: 10.1128/JB.187.5.1866-1874.2005.

Šmajs, D., Strouhal, M., and Knauf, S. (2018). Genetics of human and animal uncultivable treponemal pathogens. Infect Genet Evol 61, 92–107. doi: 10.1016/j.meegid.2018.03.015.

Staudová, B., Strouhal, M., Zobaníková, M., Cejková, D., Fulton, L. L., Chen, L., et al. (2014). Whole genome sequence of the Treponema pallidum subsp. endemicum strain Bosnia A: the genome is related to yaws treponemes but contains few loci similar to syphilis treponemes. PLoS Negl Trop Dis 8, e3261. doi: 10.1371/journal.pntd.0003261.

Sutton, M. Y., Liu, H., Steiner, B., Pillay, A., Mickey, T., Finelli, L., et al. (2001). Molecular subtyping of Treponema pallidum in an Arizona County with increasing syphilis morbidity: use of specimens from ulcers and blood. J Infect Dis 183, 1601–1606. doi: 10.1086/320698.

Taouk, M. L., Taiaroa, G., Pasricha, S., Herman, S., Chow, E. P. F., Azzatto, F., et al. (2022). Characterisation of Treponema pallidum lineages within the contemporary syphilis outbreak in Australia: a genomic epidemiological analysis. Lancet Microbe 3, e417–e426. doi: 10.1016/S2666-5247(22)00035-0.

Teufel, F., Almagro Armenteros, J. J., Johansen, A. R., Gíslason, M. H., Pihl, S. I., Tsirigos, K. D., et al. (2022). SignalP 6.0 predicts all five types of signal peptides using protein language models. Nat Biotechnol. doi: 10.1038/s41587-021-01156-3.

Thurlow, C. M., Joseph, S. J., Ganova-Raeva, L., Katz, S. S., Pereira, L., Chen, C., et al. (2022). Selective Whole-Genome Amplification as a Tool to Enrich Specimens with Low Treponema pallidum Genomic DNA Copies for Whole-Genome Sequencing. mSphere, e0000922. doi: 10.1128/msphere.00009-22.

Van Damme, K., Behets, F., Ravelomanana, N., Godornes, C., Khan, M., Randrianasolo, B., et al. (2009). Evaluation of Azithromycin Resistance in Treponema pallidum Specimens From Madagascar. Sexually Transmitted Diseases 36, 775–776. doi: 10.1097/OLQ.0b013e3181bd11dd.

Varadi, M., Anyango, S., Deshpande, M., Nair, S., Natassia, C., Yordanova, G., et al. (2022). lphaFold Protein Structure Database: massively expanding the structural coverage of protein-sequence space with high-accuracy models. Nucleic Acids Res 50, D439–D444. doi: 10.1093/nar/gkab1061.

Wang, L.-G., Lam, T. T.-Y., Xu, S., Dai, Z., Zhou, L., Feng, T., et al. (2020). Treeio: An R Package for Phylogenetic Tree Input and Output with Richly Annotated and Associated Data. Molecular Biology and Evolution 37, 599–603. doi: 10.1093/molbev/msz240.

Wick, R. R., Judd, L. M., Gorrie, C. L., and Holt, K. E. (2017). Completing bacterial genome assemblies with multiplex MinION sequencing. Microbial Genomics 3. doi: 10.1099/mgen.0.000132.

Wick, R. R., Judd, L. M., and Holt, K. E. (2019). Performance of neural network basecalling tools for Oxford Nanopore sequencing. Genome Biol 20, 129. doi: 10.1186/s13059-019-1727-y.

Wickham, H. (2016). ggplot2: Elegant Graphics for Data Analysis. 2nd ed. 2016. Cham: Springer International Publishing : Imprint: Springer doi: 10.1007/978-3-319-24277-4.

Yahashiri, A., Jorgenson, M. A., and Weiss, D. S. (2017). The SPOR Domain, a Widely Conserved Peptidoglycan Binding Domain That Targets Proteins to the Site of Cell Division. J Bacteriol 199, e00118–17. doi: 10.1128/JB.00118-17.

Yang, J., Anishchenko, I., Park, H., Peng, Z., Ovchinnikov, S., and Baker, D. (2020). Improved protein structure prediction using predicted interresidue orientations. Proc Natl Acad Sci U S A 117, 1496–1503. doi: 10.1073/pnas.1914677117.

Yu, G., Smith, D. K., Zhu, H., Guan, Y., and Lam, T. T. (2017). ggtree: an R package for visualization and annotation of phylogenetic trees with their covariates and other associated data. Methods Ecol Evol 8, 28–36. doi: 10.1111/2041-210X.12628.

Yu, N. Y., Wagner, J. R., Laird, M. R., Melli, G., Rey, S., Lo, R., et al. (2010). PSORTb 3.0: improved protein subcellular localization prediction with refined localization subcategories and predictive capabilities for all prokaryotes. Bioinformatics 26, 1608–1615. doi: 10.1093/bioinformatics/btq249.

Zhou, L., Feng, T., Xu, S., Gao, F., Lam, T. T., Wang, Q., et al. (2022). ggmsa: a visual exploration tool for multiple sequence alignment and associated data. Briefings in Bioinformatics, bbac222. doi: 10.1093/bib/bbac222.

Zobaníková, M., Strouhal, M., Mikalová, L., Cejková, D., Ambrožová, L., Pospíšilová, P., et al. (2013). Whole genome sequence of the Treponema Fribourg-Blanc: unspecified simian isolate is highly similar to the yaws subspecies. PLoS Negl Trop Dis 7, e2172. doi: 10.1371/journal.pntd.0002172.

