## Supplementary figures and images for "High-throughput nanopore sequencing of *Treponema pallidum* tandem repeat genes *arp* and *tp0470* reveals clade-specific patterns and recapitulates global whole genome phylogeny"

### Supplementary Figure 1

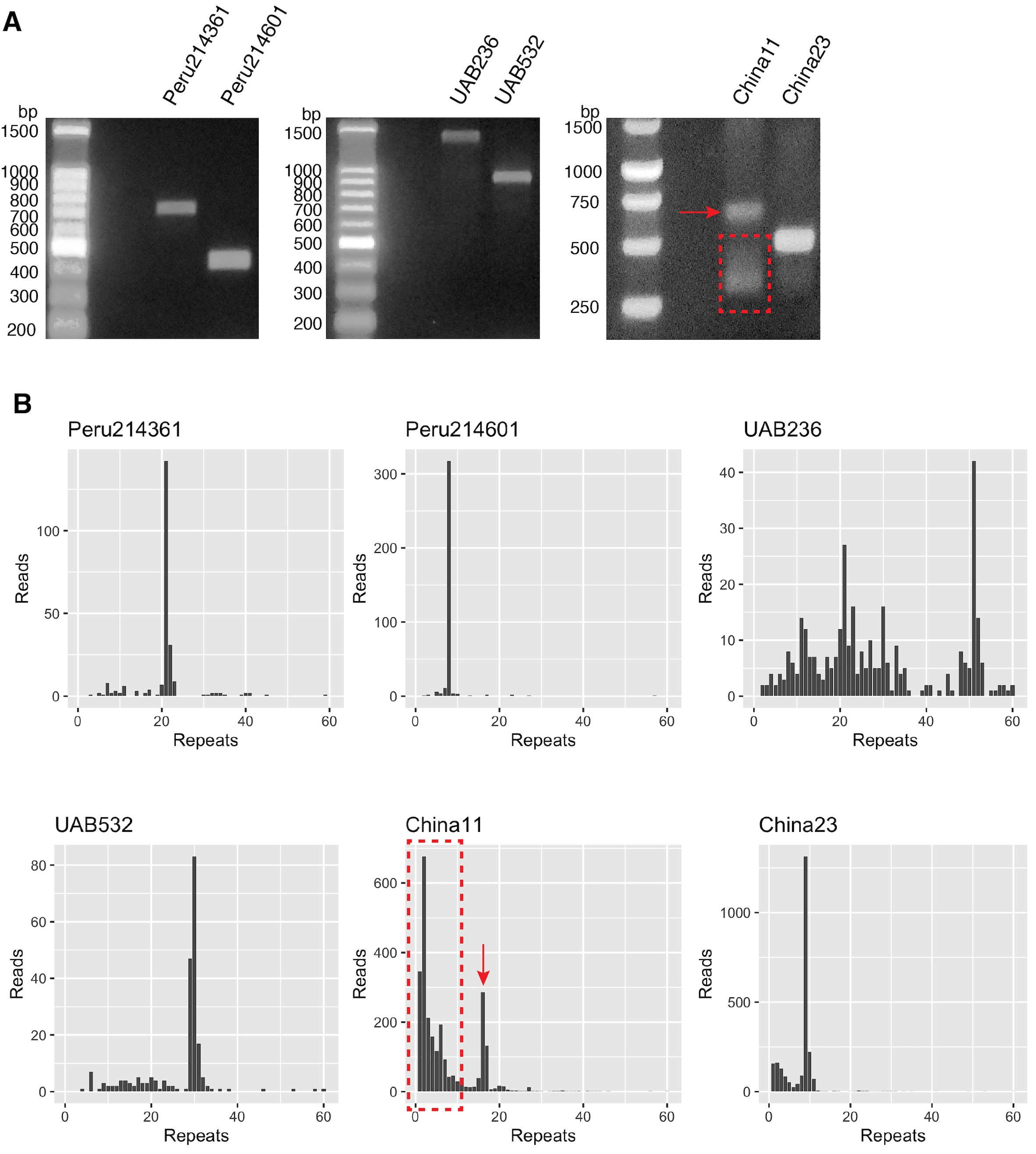

### Supplementary Figure 2

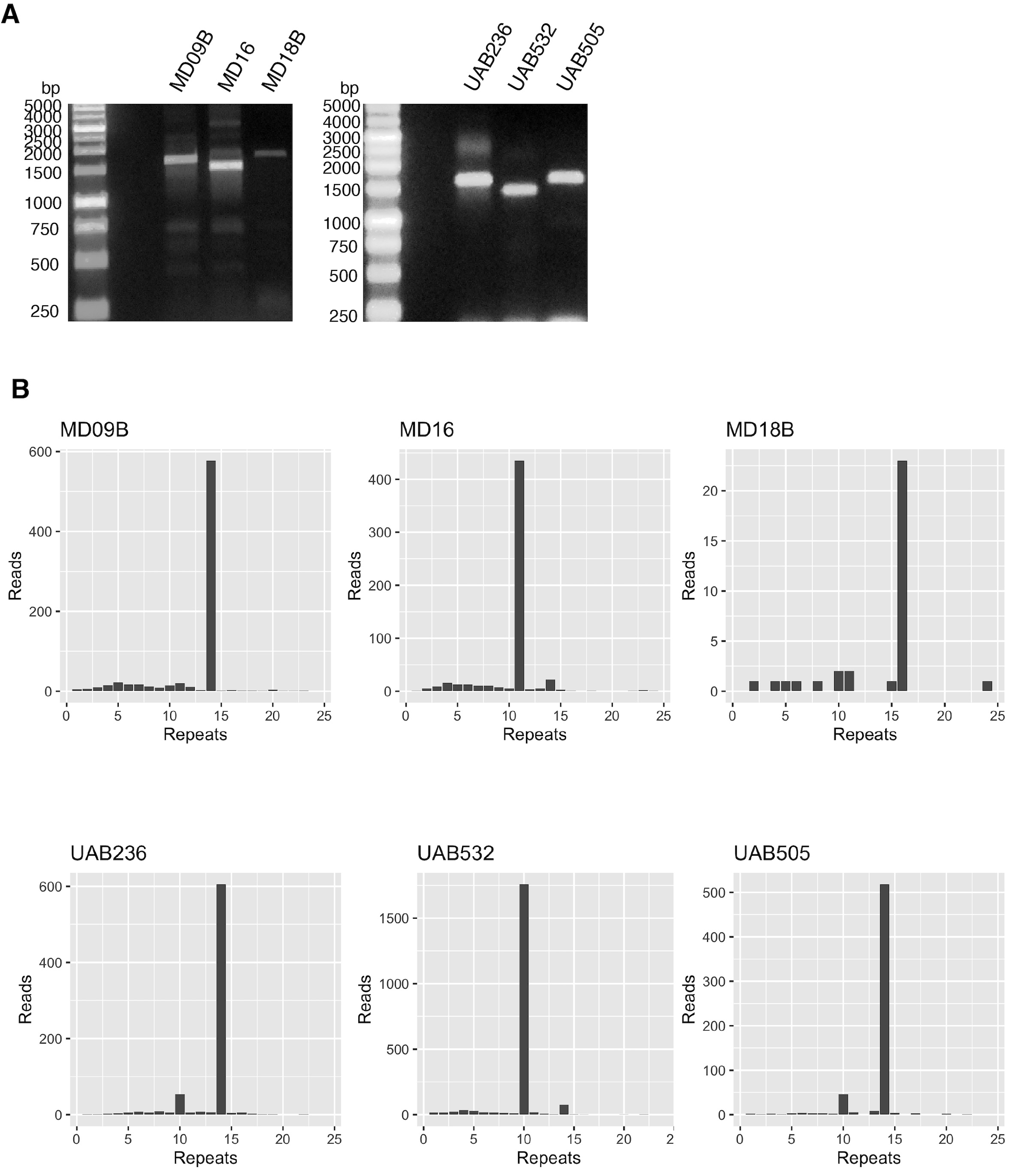

### Supplementary Figure 3

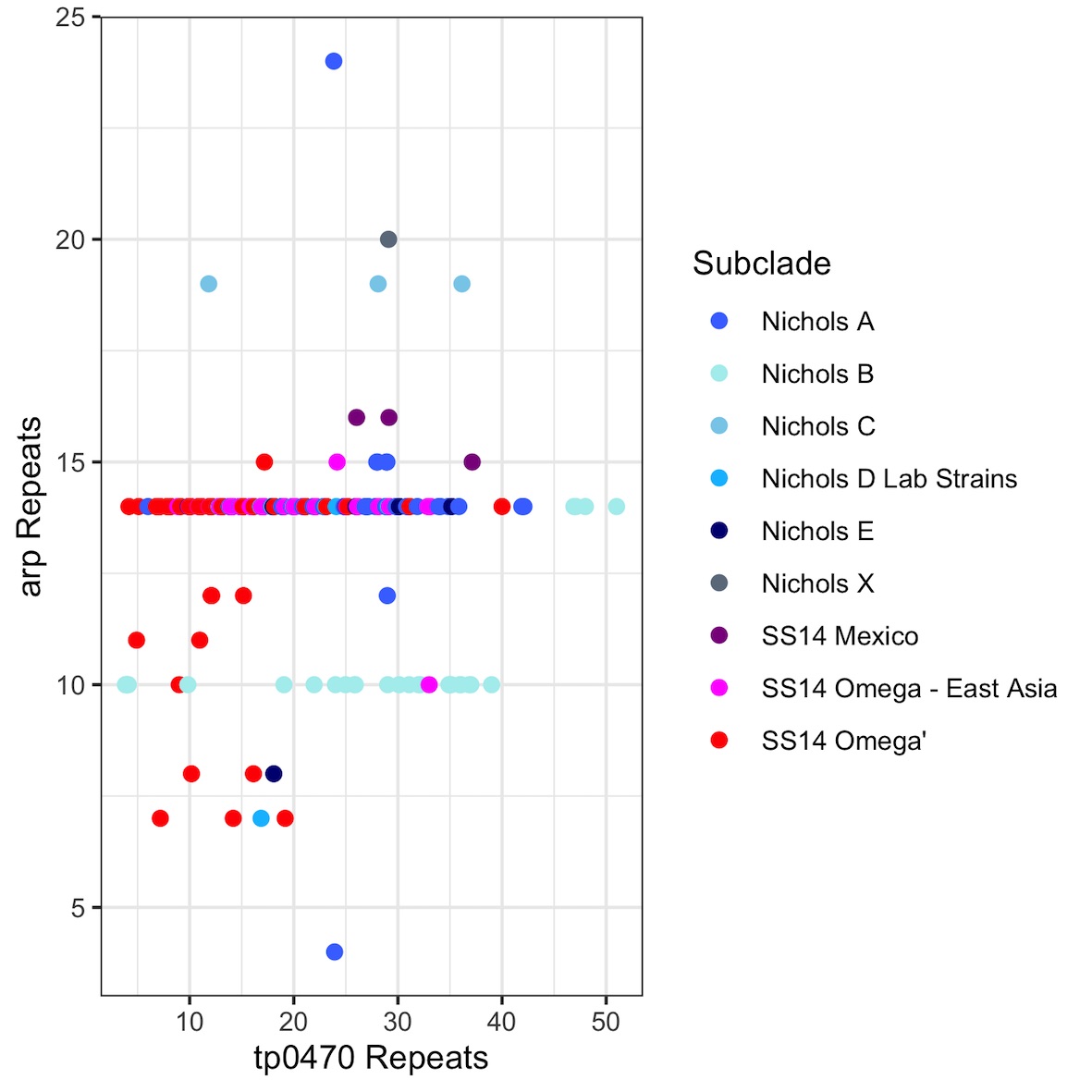

### Supplementary Figure 4

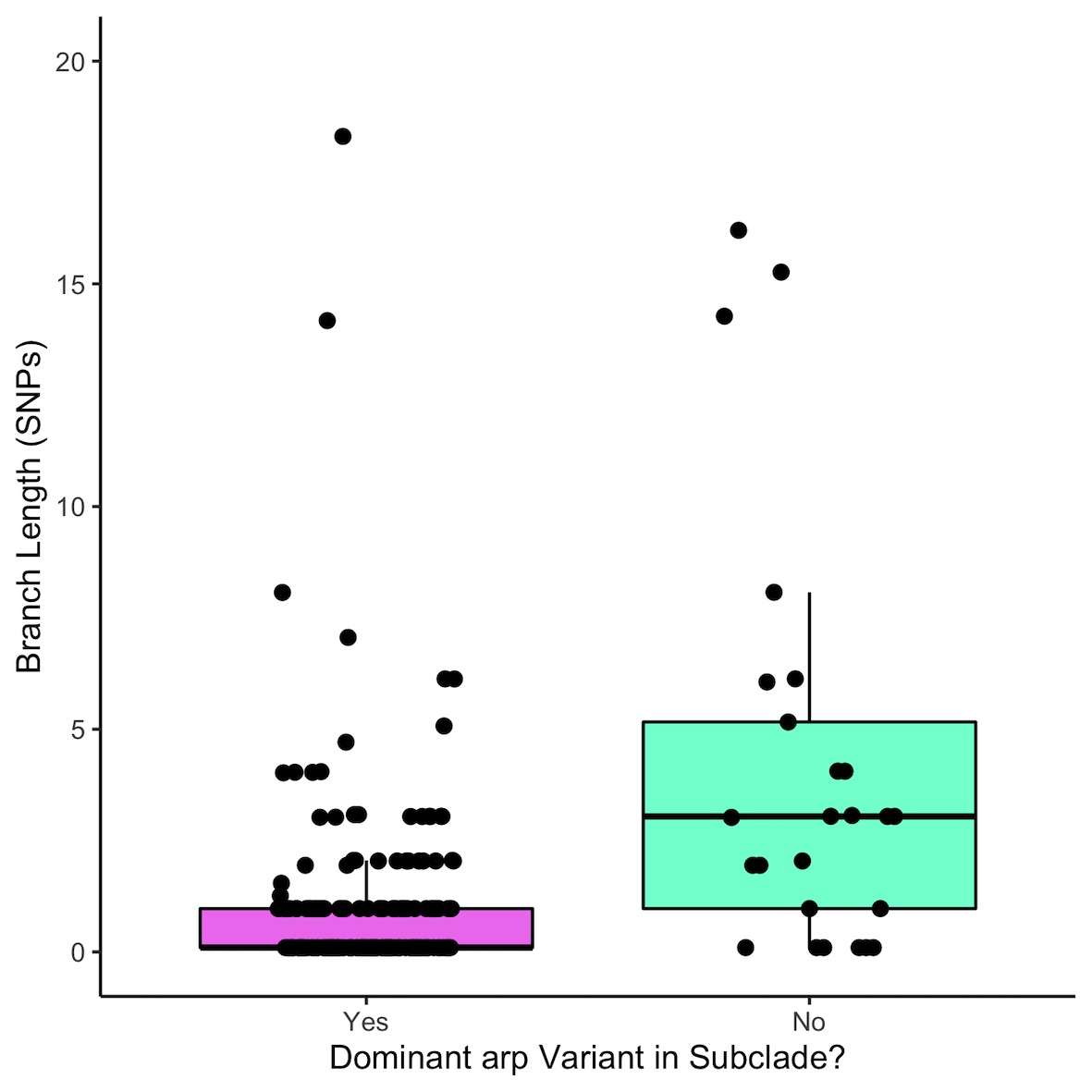

### Supplementary Figure 5

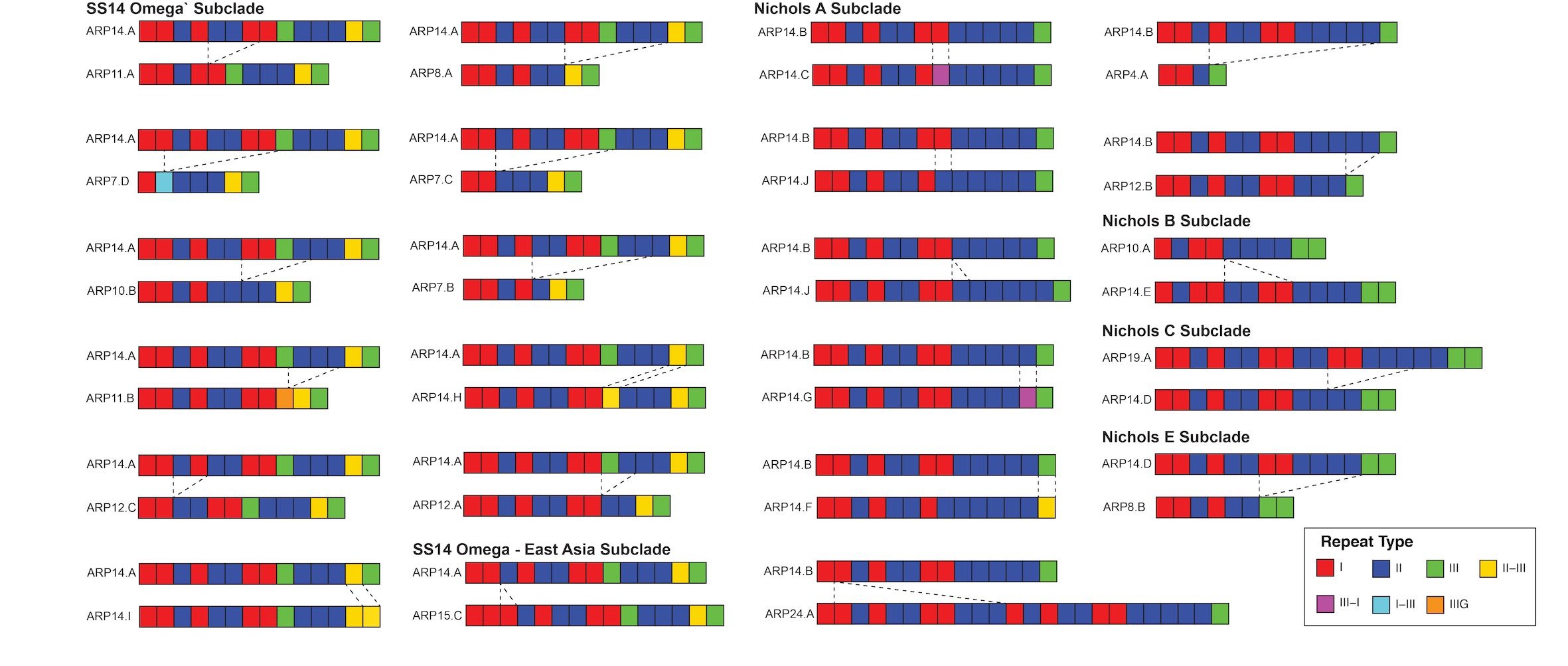

### Supplementary Figure 6

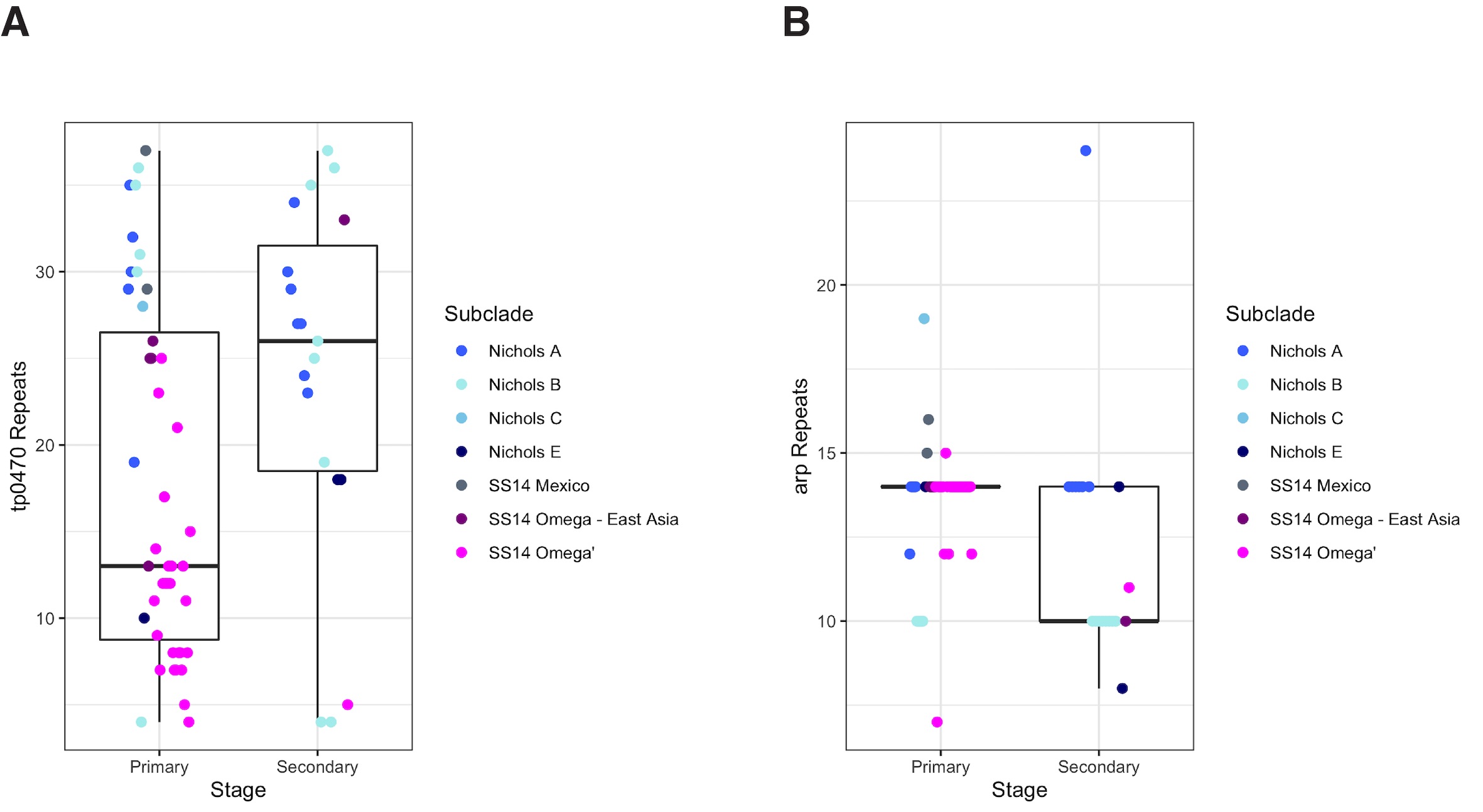

### Supplementary Figure 7

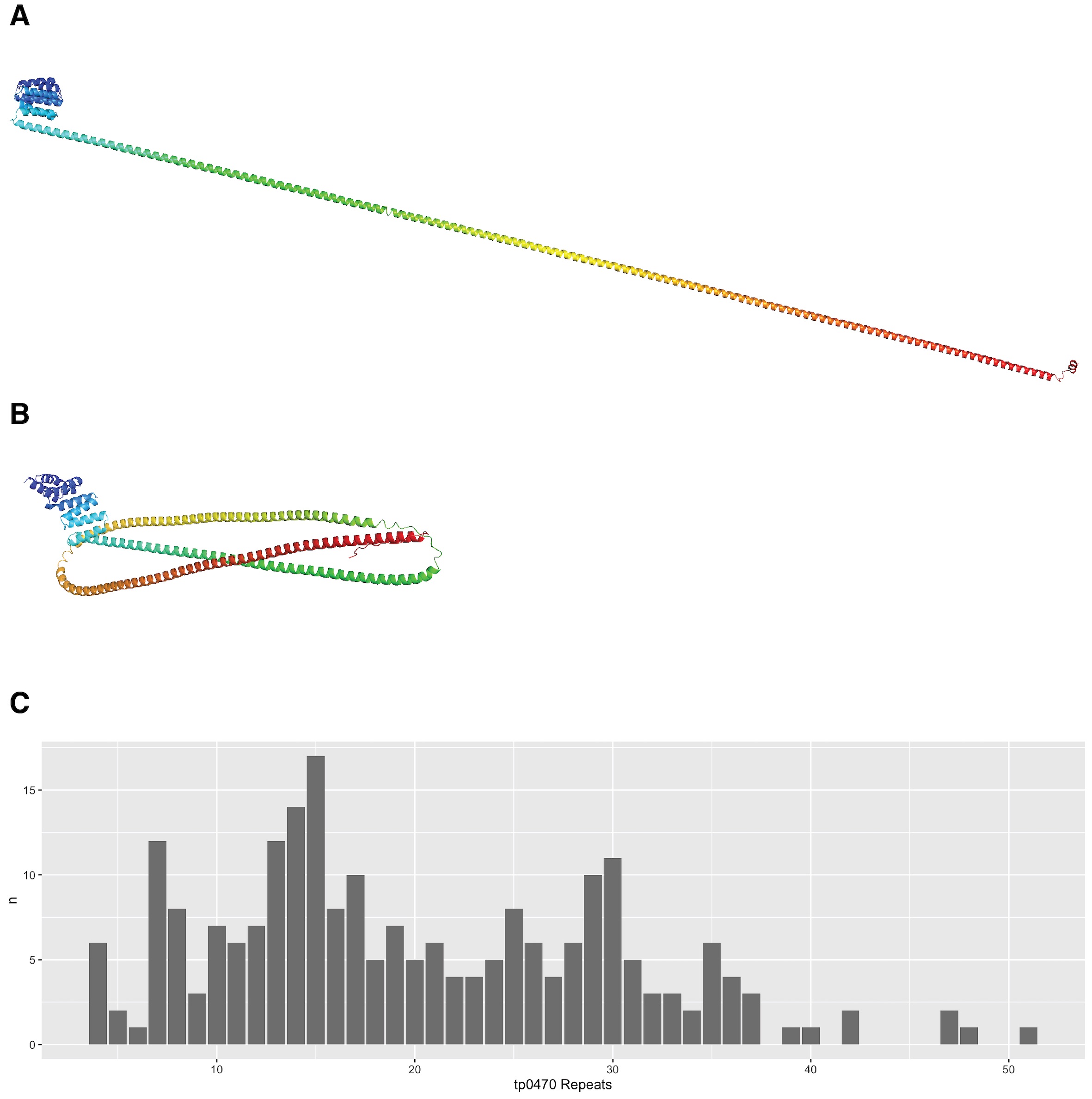

### Supplementary Figure 8

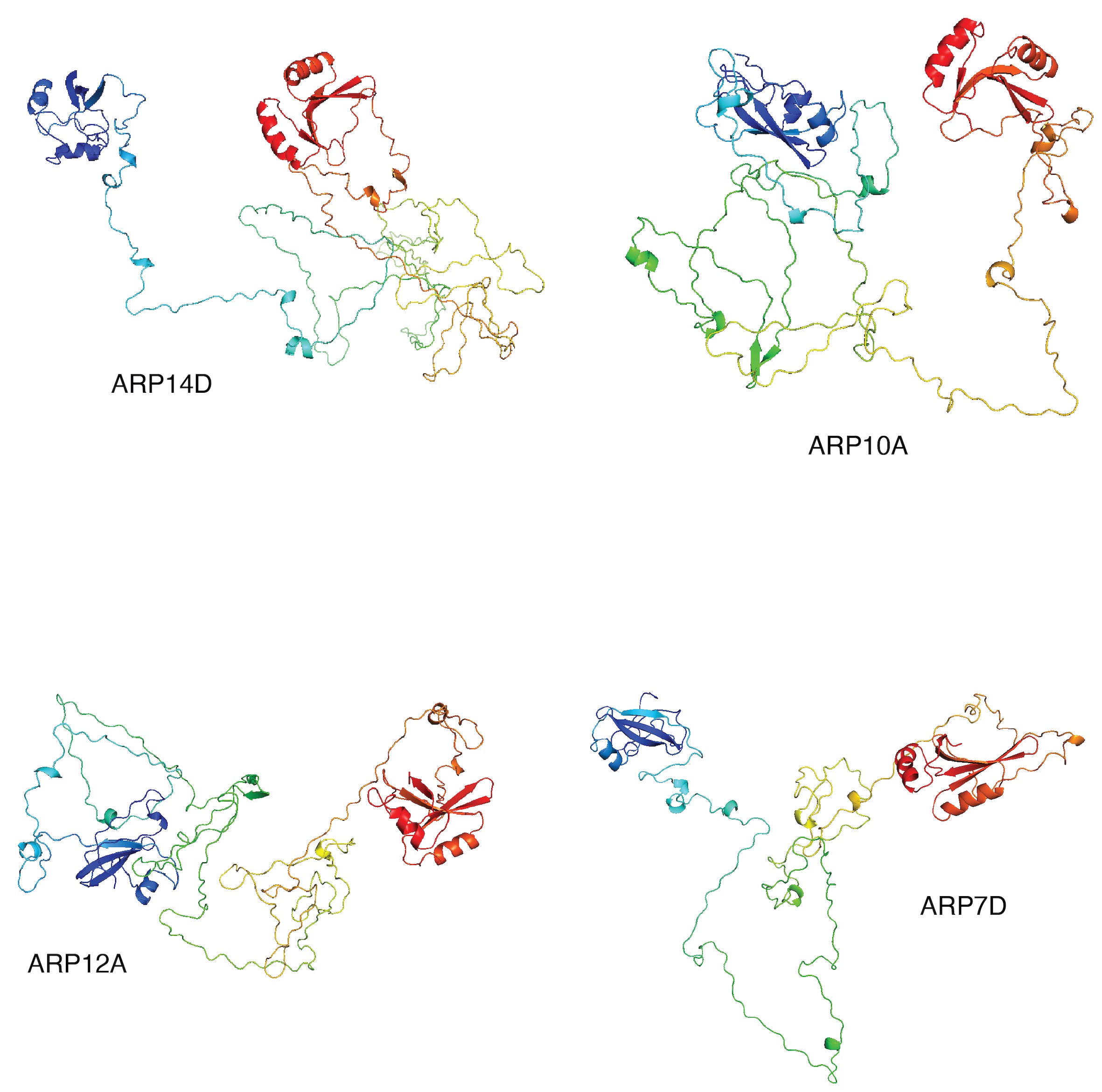
